# Validating the Representational Space of Deep Reinforcement Learning Models of Behavior with Neural Data

**DOI:** 10.1101/2021.06.15.448556

**Authors:** Sebastian Bruch, Patrick McClure, Jingfeng Zhou, Geoffrey Schoenbaum, Francisco Pereira

## Abstract

Deep Reinforcement Learning (Deep RL) agents have in recent years emerged as successful models of animal behavior in a variety of complex learning tasks, as exemplified by Song et al. [2017]. As agents are typically trained to mimic an animal subject, the emphasis in past studies on behavior as a means of evaluating the fitness of models to experimental data is only natural. But the true power of Deep RL agents lies in their ability to learn neural computations and codes that generate a particular behavior—factors that are also of great relevance and interest to computational neuroscience. On that basis, we believe that model evaluation should include an examination of neural representations and validation against neural recordings from animal subjects. In this paper, we introduce a procedure to test hypotheses about the relationship between internal representations of Deep RL agents and those in animal neural recordings. Taking a sequential learning task as a running example, we apply our method and show that the geometry of representations learnt by artificial agents is similar to that of the biological subjects’, and that such similarities are driven by shared information in some latent space. Our method is applicable to any Deep RL agent that learns a Markov Decision Process, and as such enables researchers to assess the suitability of more advanced Deep Learning modules, or map hierarchies of representations to different parts of a circuit in the brain, and help shed light on their function. To demonstrate that point, we conduct an ablation study to deduce that, in the sequential task under consideration, temporal information plays a key role in molding a correct representation of the task.

## 1 Introduction

For decades Reinforcement Learning (RL) [Sutton and Barto, 2018] has proven instrumental in our understanding of the neural mechanisms of reward-driven learning and decision making by way of providing a formal language to explain a wealth of empirical phenomena [Niv, 2009]. The scope and complexity of what this framework could formalize broadened dramatically when, in recent years, researchers coupled RL with Deep Learning [Goodfellow et al., 2016]—giving rise to what is commonly referred to as “Deep RL” [Botvinick et al., 2020, Dabney et al., 2020, Song et al., 2017, Wang et al., 2018]. For example, Wang et al. [Wang et al., 2018] showed how an RL model that uses recurrent neural networks (RNN) [Hochreiter and Schmidhuber, 1997] helps explain *meta learning* in the prefrontal cortex. Dabney et al. [Dabney et al., 2020], as another example, extended the notion of dopamine-driven value learning from *expected* reward to reward *distribution*, to help explain how the brain manages uncertainty in the reward distribution. Across studies, it is compelling to witness that it is possible to have a Deep RL agent learn to perform a task given only the same stimuli and reinforcement schedule as an experimental subject.

For all the enthusiasm surrounding Deep RL in the neuroscientific literature, however, existing studies seldom look beyond behavior for evaluation, with few exceptions [Bernardi et al., 2020, Dabney et al., 2020, Kao, 2019, Yang et al., 2017, Zhang et al., 2018, 2020]. This focus on whether or not models correctly predict an observed behavior means that much of the literature, when formulating or evaluating a model, ignores the factors that *generate* that behaviour. That approach, as we argue below, is misguided, wasteful, and insufficient.

It is misguided because models as complex as Deep RL agents can produce arbitrary behaviors with trivial adjustments. Simply arriving at a particular behavior that happens to correlate with that of a biological subject’s, therefore, says nothing about the similarity or dissimilarity of the neural mechanisms that generated that behavior. It is wasteful because it ignores the often rich representations that Deep Learning models— more precisely Deep Neural Networks (DNNs)—are known to learn. That power to learn latent neural codes for perceptual observations and hidden structures, after all, is what has fueled great strides on myriad scientific fronts such as natural language processing, computer vision, and reinforcement learning (c.f., Botvinick et al. [2020] and references therein). By not examining the learnt representations and investigating their potential link to neural recordings obtained from the brain, one risks discarding valuable information.

Finally, a narrow focus on behavior alone is insufficient because neural computations and codes are, in many cases, of greater interest than the behavior to which they lead. Consider, for instance, one of the more prominent hypotheses on the function of the orbitofrontal cortex (OFC), which argues that the OFC is part of a circuit that learns a “cognitive map” of the task space [Behrens et al., 2018, Constantinescu et al., 2016, Garvert et al., 2017, Schuck et al., 2016, Wilson et al., 2014, Zhou et al., 2019, 2021]. That hypothesis, by its very definition, rests on the notion of the formation and evolution of neural representations.

For the reasons above, it is encouraging to witness a shift, albeit a still-inchoate one, towards more in-depth analyses of Deep RL agents [Kao, 2019, Yang et al., 2017, Zhang et al., 2018, 2020], the focus on qualitative rather than quantitative assessment notwithstanding. One study by Bernardi et al. [Bernardi et al., 2020] stands out as the authors attempt to characterize *abstraction* quantitatively: Observing that neural representations in the brain of monkeys encode latent variables of a reversal-learning task, the authors test how accurately a linear readout of variables from neural recordings generalizes to the unseen portion of the data. An accuracy bounded away from random chance is taken to suggest that the brain has learnt an abstract representation of the task features.

While encouraging, these incipient ideas do not yet offer a systematic way of testing hypotheses about neural representations and examining the internal representations in Deep RL agents. Few studies [Zhou et al., 2019, 2021] address inherent challenges in analyzing neural codes such as manifold misalignment due to inconsistencies in the number of recorded neurons, among other factors. None to date, to the best of our knowledge, directly compares artificially learnt representations with biological ones for reinforcement learning problems. Stating and testing hypotheses on the space of neural representations are largely absent from the literature and remain a black art.

In this work, we set out to address the points above. Our goal is to show that, by raising a series of targeted questions, it is easy to reason about the spaces of neural representations in general. In particular, given two or more disparate sets of neural representations—obtained from machines, subjects, or both—we investigate (a) whether they are geometrically similar, (b) whether an observed geometrical similarity is driven by common latent factors, and (c) whether those common factors encode similar information.

These research questions probe what *causes* behavioral similarities and help us shift away from analyzing just the symptoms. In other words, if a Deep RL model behaves the same way as a biological subject on a particular task, the questions we raise here help to determine what factors, if any, contribute to that similarity. These factors can be viewed as the constructs one ultimately wants to discover, identified via modelling of task performance and validated in neural data.

As we unpack these research questions, we discuss specific technical challenges. We show, for example, how a combination of standard dimensionality reduction and manifold alignment techniques facilitates a direct comparison of otherwise incompatible sets of representations. Building on Representational Similarity Analysis [Kriegeskorte et al., 2008, Popal et al., 2020] and utilizing non-parametric statistical tests, we show how one may assess significance of measured similarities between representational spaces. Finally, as a testament to the generality of this method, we compare the representational space of an artificial agent with a biological subject, and through an ablation study, demonstrate how our methods can be used to decide which Deep Learning components lead to more accurate models of neural data.

Throughout this work, we ground our discussion in the learning task of Zhou et al. [2021], stating and testing hypotheses in that context. In that study, rats learn to solve an odor sequence problem and generalize their knowledge to structurally-similar but perceptually-different problems. During training, the authors record different sets of OFC neurons on different days, thereby capturing neural representations from the population. As one of the objectives of our study is to compare models with subjects, we train a Deep RL agent with an RNN on the same task and record its internal representations. We then apply our analysis to the two sets of representations.

We note that our method is agnostic to the specific task at hand and we only choose to focus on a particular task for concreteness. We chose this particular task for two reasons. First, as already alluded to, neural representations are key to examining the cognitive map hypothesis. Second, as the focus of the task is on abstraction and generalization from one problem to another, it gives us the opportunity to apply our method to compare neural recordings from one problem with recordings from another, both obtained from the same set of biological subjects. That last point means that, in the end, we can use our method to compare biological neural representational spaces not just with artificial ones but also with other biological spaces, thereby establishing a baseline.

The remainder of this paper is organized as follows. We begin, in Preliminaries, with a short overview of the sequence learning task [Zhou et al., 2021] and a description of a simple Deep RL agent that will be used as a possible model of the OFC in the context of an odor sequence problem. Methodology provides a detailed introduction to the three research questions that comprise our method, and the motivation for each. In Results, we apply our method to the datasets of neural recordings from Zhou et al. [2021] and our artificial agents. We discuss these results and the scope of applicability of the method in Discussion.

## 2 Preliminaries

As noted earlier, we tie the presentation of our methodology to the work of Zhou et al. [2021], stating and testing hypotheses for their task for concreteness. To give context, then, we briefly describe the task itself and the neural data acquired in that study in The Odor Sequence Problem. Moreover, our methodology requires neural network-based artificial agents to serve as models of animal behaviour, in the manner introduced in Song et al. [2017]. We developed two Deep RL agents for the odor sequence task, similar to those in past studies [Song et al., 2017, Wang et al., 2018] that have shown to be reasonable models of the prefrontal cortex. These are described in Artificial Agent.

### 2.1 The Odor Sequence Problem

The goal of the “odor sequence” task in Zhou et al. [2021], broadly, was to study the role of the orbitfrontal cortex (OFC) in abstraction and generalization. The research question there concerned how rats learn to solve a complex problem and generalize what they learn to other structurally-similar problems. We review the design of their task in this section.

The odor sequence task, illustrated in Figure 1 (Right), is a series of five independent “problems,” labeled *A* through *E*. Subjects spend 15 days on one problem and subsequently proceed to the next one in alphabetical order. These problems are instances of a common template, all sharing the same task structure but each using a distinct set of odors as stimuli. It suffices then to describe a single problem.

**Figure 1:**
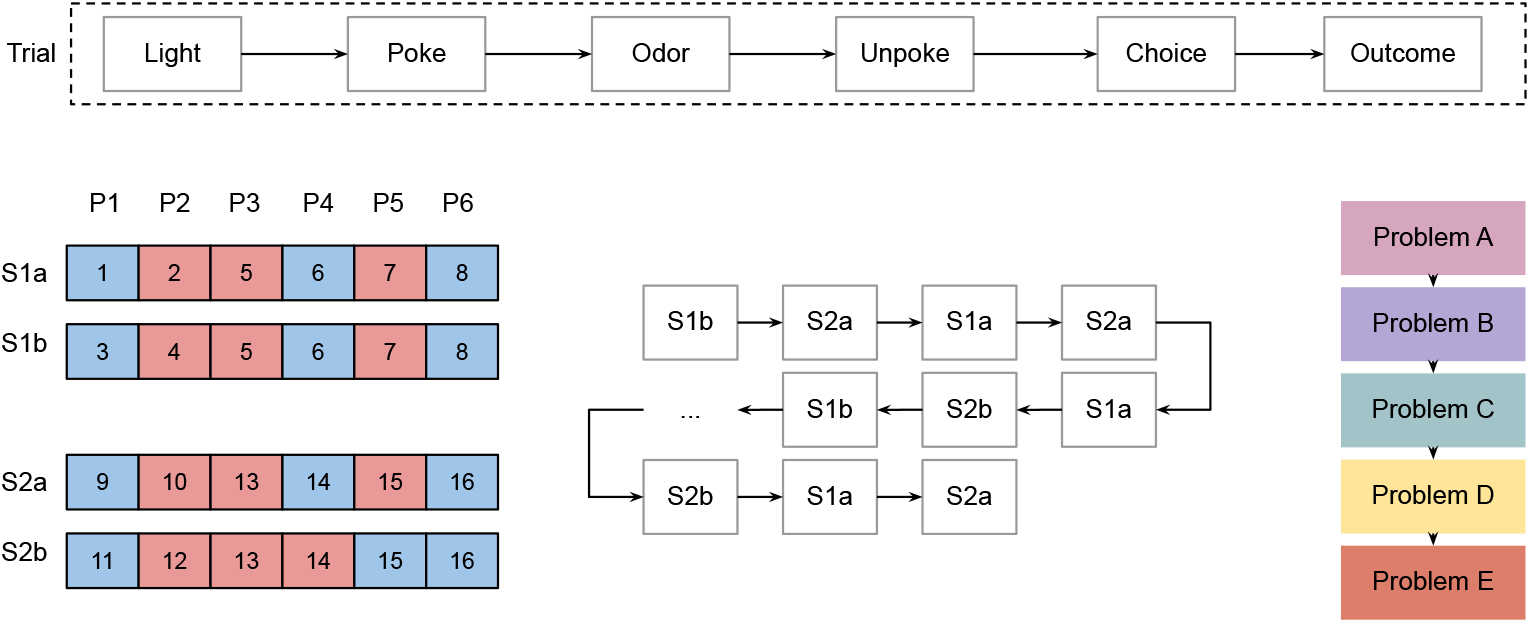
Illustration of the odor sequence task [Zhou et al., 2021]. **Top**: A trial is made up of a number of events: “Light” event to signal that the animal should go to the odor port; “Poke” event occurs when the animal enters the odor port; “Odor” event is when a trial-specific odor is released; “Unpoke” and “Choice” events when the animal leaves the odor port and makes a decision; and, finally, “Outcome” is when the outcome of the trial is revealed to the animal. **Left**: Trials (individual squares) are annotated with their odor identifier (1–16). Six trials put together in positions “P1” through “P6” make up a “sequence.” Whether a trial is rewarded (blue) or not (red) is a function of the sequence it is in and its position within the sequence. The reward distribution is the same in all four sequences with the exception of positions P4 and P5 in sequence S2b. **Center**: A series of 80 sequences alternating between S1 and S2 result in 480 trials, which we call a “problem.” **Right**: The task is a series of five problems, where each problem has its own unique set of 16 odors (i.e., odor 1 in Problem *A* is different from odor 1 in all other problems). The animals spend 15 days on each problem.

A problem is a succession of 80 “sequences.” It is constructed by a repeated arrangement of four unique sequences—S1={S1a, S1b} and S2={S2a, S2b}—that alternate between S1 and S2; in the end, there are 20 occurrences of each, as illustrated in Figure 1 (Center). A sequence itself is made up of 6 positions, where each position is, deterministically, either “rewarded” or not. Sequences S1a, S1b, and S2a share the same reward distribution, whereas in S2b, the rewards for positions 4 and 5 are reversed, as shown in Figure 1 (Left).

Finally, in each position of every sequence a “trial” takes place—that, then, results in 24 unique trials. In a trial, a light cues the dispensing of an odor that has been prescribed for that position and sequence. Upon the light cue (“light” event), the subject must move to the odor port (“poke” event), must receive the odor (“odor” event), and must leave the port (“unpoke” event) to ultimately choose to visit a sucrose well or not—whether a reward is found in the well depends on the reward status of the current position. This is depicted in Figure 1 (Top).

### 2.2 Neural Data

During a fixed time period before and after every event in every trial (e.g., light, poke, odor, unpoke), single-unit neuronal activity in the subjects’ OFC is recorded. Pre-event time is set to 200ms and the post-event time to 600ms, with spike counts averaged within 100ms bins. That results in 8 time points per event per neuron. We highlight that, the set of recorded neurons in each animal is not consistent over days and problems. We refer the reader to Zhou et al. [2021] for a more comprehensive description and a technical discussion of the neural recordings.

Zhou et al. [Zhou et al., 2021] found that as subjects navigate this complex task—from trial to trial, sequence to sequence, and problem to problem, all as a continuum—they learn the reward distribution and discover the problem template. That such structure discovery is noteworthy is because it happens despite the fact that the perceptual signals— the odors—vary between problems. More notable still is the evolution of neuronal activity in the OFC: As the days progress, OFC neurons become more selective, leading to more refined firing patterns that, in effect, come to distill the knowledge required to solve this sequential task and to generalize to other structurally similar tasks.

Throughout this work, we take as the neural representation of an *event* the firing rate of all recorded neurons, through all 8 time points, concatenated into a single vector of values. By putting together these values from the light, poke, odor, and unpoke events, we arrive at the neural representation of a *trial.* That choice mirrors the method of Zhou et al. [2021]. We elaborate the precise construction of neural representations in Methodology. Finally, as we are interested in the outcome of learning, rather than its evolution, we generally limit our analysis to the OFC representations from the final day of training (i.e., day 15) on each problem unless indicated otherwise.

### 2.3 Artificial Agent

We use a standard neural network-based reinforcement learning agent that has proven particularly effective at modeling a number of learning tasks in the prefrontal cortex [Song et al., 2017, Wang et al., 2018]. For reasons that will become apparent shortly, we refer to this agent as A3cRnn. In this section, we describe the agent’s input, architecture, output, and training procedure. We will also present an ablated variant, dubbed A3cFfn, that lacks the ability to model temporal signals. But, to give context to our discussion, we begin with a description of the simulated task.

#### 2.3.1 Simulated Odor Sequence Problem

The task on which we train our reinforcement learning agents is a reduction of the odor sequence problem. The number of problems and the structure of sequences remain as before, but what takes place within a trial and the array of actions available to the agent take a different form. In particular, a simulated trial consists only of two events, light and odor, and the agent may only choose between three actions: going to the odor port, going to the sucrose well, and staying idle.

In the *light* event, we present to the agent the light cue, an identifier that is the same across all problems. The correct action in this case is to “go to the odor port.” All other actions result in a reward of —1 and an immediate transition to the next trial. In the event the agent takes the correct action, it enters the *odor* event but receives no reward (i.e., a reward of 0).

In the *odor* event of a trial, the agent receives an odor identifier whose value depends on the position and sequence of the trial—there are 16 distinct odors in each problem. The sets of odors are disjoint between problems, resulting in a total of 80 identifiers representing all five sets of odors. This arrangement rests on the assumption that the animals are able to differentiate all 80 odors. There are two meaningful actions once the agent experiences an odor: A positive action to collect its reward or a negative action to stay idle. The agent earns a reward of +1 if it takes the positive action in a rewarded trial, no reward if it takes the negative action in a non-rewarded trial, and a reward of — 1 in all other cases, including when the agent mistakenly chooses the “go to the odor port” action.

We should note that, the adopted discretization of the space of events and actions, while greatly simplifying the problem setup and minimizing the run-time of our experiments, ultimately does not affect our analysis. For example, even though we included the light event in the simulated task, the agent quickly learns to disregard it as irrelevant to the objective of the task, and learns a trivial mapping from “light” (state) to “going to the odor port” (policy). We observe a similar pattern with the addition of other event types that are inconsequential to the outcome and can be safely ignored by the agent. On the other hand, the odor event, as in the real experiments, contains the factors that lead up to the decision.

#### 2.3.2 Input to the Agent

At each of the two events, the agent observes an identifier which either represents “light” in the light event or an “odor” in the odor event. In order to present such observations to the agent, we encode their identifiers as a categorical variable that can take one of 81 distinct values. In particular, we employ a *one-hot encoding* scheme where each identifier is a binary vector of 81 elements with only one set element. For example, light would be represented as (1,0,..., 0), the first odor of the first problem as (0,1,0,..., 0), and so on.

In addition to the observation above, we explicitly make the chosen action and its reward outcome from the preceding event available to the agent as input. Actions are represented using one-hot encoding as a vector of three binary values, one slot for every available action. Reward, on the other hand, is represented as an integer. These input features together form the input layer in our model.

#### 2.3.3 Agent Architecture and Training

Before we delve into the details of the models used in this work, it is important to note that we are less interested in modeling per se than we are in validation. In other words, we are not concerned with the specific choices of Deep Learning components and their hyperparameters. Instead, we assume there exists a pool of models to choose from, and propose a method to evaluate their validity purely based on neural representations.

Following the reasoning above, we adopt an agent architecture drawn from prior work [Song et al., 2017, Wang et al., 2018] that is effective in modeling sequential tasks from a behavioral standpoint. We then perform an ablation to create a second architecture, and contrast these two candidates in the remainder of this article. Our choice of hyperparameters is also guided by previous work [Song et al., 2017] with a limited fine-tuning effort aimed at reducing the capacity of the agents while still ensuring that the agents behave reasonably. We must, however, note that the method we present here can be used just as easily to search through any other extension or ablation, or other choices in the space of architectures and hyperparameters.

With the note above in mind, we are ready to present the specifics of our agents. We use a feed-forward neural network to encode the input to the model. This network consists of three layers with 256, 128, and 64 artificial neurons respectively. Each neuron uses the Rectified Linear Unit (ReLU) activation function, and is fully connected to the neurons in the next layer.

We compose the input encoder above with a function whose role, we hypothesize, is to emulate the OFC, and whose activity is the analogue of neural data. This function, in fact, is the only component that is different between the agents: In A3cRnn, it is a Gated Recurrent Unit [Cho et al., 2014], a specific type of the more general Recurrent Neural Network (RNN), whereas in A3cFfn, it is a single layer of ReLU-activated neurons. This small architectural difference between the two agents manifests in the way the two learn to solve the same task: A3cRnn reuses certain neural codes from previous time steps to arrive at a decision in the current time step, thereby injecting its knowledge of past states directly into learning and decision making. In contrast, A3cFfn has no such mechanism to incorporate information about preceding states. A comparison of these two models should, therefore, highlight a fundamental aspect of the function of the OFC. In both agents, the number of hidden neurons is 32. Figure 2 shows the model architecture for A3cRnn and highlights its difference with A3cFfn.

**Figure 2:**
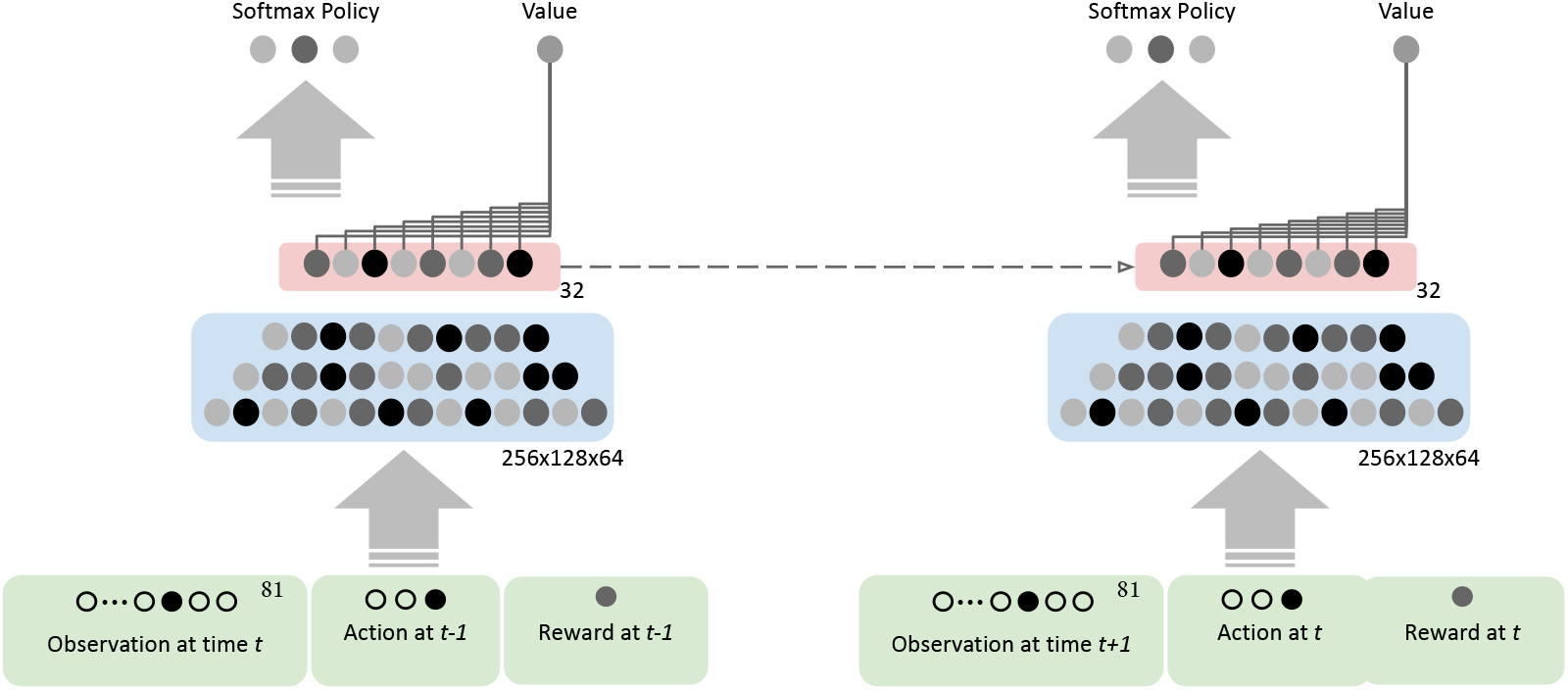
A3cRnn model architecture. (**Left**) The input layer (bottom, green boxes) is made up of the following: The current observation at time *t* which is either the code for “light” or an “odor”; action taken by the agent at the previous time step (*t* – 1); and reward given to the agent at time *t* – 1. A feed-forward network then encodes the input to the network (middle, blue box). The encoded input is then passed onto an RNN unit (top, red box). A linear readout of the output of the RNN approximates the value of the current state. The agent also learns a policy by defining the probability of taking each of the three actions to be proportional to a linear function of the output of the RNN, one per action. The agent then executes the action that has the highest probability, and receives a reward from the environment accordingly—the executed action and earned reward are fed to the model as input at the next time step. (**Right**) We unrolled the network to illustrate the recurrent nature of the model, where the output of the RNN at time t is used as input to itself at *t* + 1. This is highlighted by the dashed line connecting the RNN module to itself at the next time step. Note that, by removing this recurrent connection, the model reduces to A3cFfn.

The output layer, much like the input layers, is common between the two agents. It consists of value and policy networks. The value network is a linear function whose output is an approximation of the value of the current state. The policy network consists of three linear functions (one per action), that together, using a softmax mapping, determine a probability distribution over actions: The agent takes the action with the largest probability.

We train the agents using the Asynchronous Advantage Actor-Critic (A3C) algorithm [Mnih et al., 2016], where the cost function under optimization is a linear combination of the policy gradient, temporal difference of the value [Sutton and Barto, 2018], and a regularizing entropy term (with weight 0.3). Gradient updates are applied every 120 steps, where one step equals the completion of one event. We use a discount factor of 0.5 when approximating returns.

Finally, during training, the agent completes 100 episodes per problem, where an episode is defined as visiting all 480 trials of a problem. We note that, deciding when to terminate the training was based on behavior: In our experiments, we observed that 100 episodes were often sufficient to guarantee convergence.

## 3 Methodology

We present a detailed description of our methodology in this section. We begin with the transformation of neural recordings—obtained from a biological or artificial source—into neural representations whereby a trial is represented by a vector of real numbers. In this way, we translate questions about neural data into questions about the geometrical properties of abstract vectors. Such a formulation allows us to remain agnostic to the source of representations, thereby facilitating comparisons of neural recordings within and across subjects and agents.

Given this abstract notion, we ask whether two sets of neural representations have similar structures: Is the degree of similarity between pairs of trials consistent across problem instances or sources? To that end, we formulate a null hypothesis in terms of representational similarities and, to test it, introduce a non-parametric statistical procedure. We then describe a procedure to test whether the structural consistency present in two or more spaces is driven by common latent factors. Finally, we lay out a procedure to test the information contained in those common latent factors. In particular, with this method we can determine whether knowing representations in one space allows us to deduce the optimal decision in another space.

### 3.1 Obtaining Neural Representations from Neural Data or Artificial Agents

Neural representations, whether obtained from biological subjects or artificial agents, form the basis of our analyses. While every dataset of neural representations is naturally only meaningful in the context of a specific task and, in that sense, is unique, we can nonetheless conceptualize every such dataset in the same way: As a function that maps an input (e.g., state, trial, or event) to a vector of real numbers. For example, in the odor sequence task described earlier, neural recordings from the OFC can be viewed as a mapping from a particular event to a vector of (normalized) activity levels recorded from the OFC as the subject experienced that event. We take this view and only require that such a mapping exist; our method is agnostic to *how* this mapping is formed.

In this section, we make the above more concrete by describing how neural recordings from the OFC or from any of the artificial agents under consideration can be stated as a function of visits to trials. This formalism simplifies the discussion in upcoming sections and, further, serves as an example that can be extended in obvious ways to arbitrary datasets of neural representations.

For brevity, let us introduce the following notation to compactly refer to the terminology reviewed earlier. We denote by 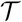 the set of 24 unique trials, {S1a1–S1a6, S1b1–S1b6, S2a1–S2a6, S2b1–S2b6}, and let t serve as an index into 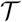. Each trial is visited 20 times through the course of a single problem. We denote a visit by 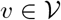 with 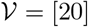 being the index set of all visits from 1 to 20. Finally, we let *p* be a particular problem, one of 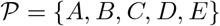.

#### 3.1.1 OFC Neural Representations

We define 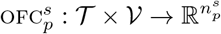 to be a mapping from visits of trials in problem 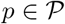 to its 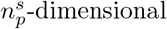 OFC representation obtained from subject *s*. Said differently, 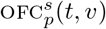 is the representation expressed in the OFC of subject *s* when it visits trial *t* for the *v*^th^ time in problem *p*. In this work, as in Zhou et al. [2021], this representation is a concatenation of a subject’s OFC neuronal activity during the light, poke, odor, and unpoke events. Note that, in general, we may not assume that 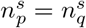 for 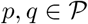 if *p* ≠ *q*.

For consistency with Zhou et al. [2021], we work at the ensemble level. That means aggregating the functions 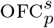 along subjects, to form one monolithic, imagined subject whose OFC is an amalgamation of the participating subjects’. Following that work, given nine subjects, we define 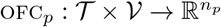 as follows:

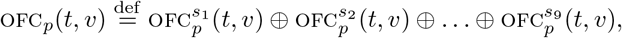

where ⊕ is the concatenation operator. It is clear that 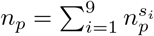 is the dimensionality of the final representation. Finally, to reduce the effect of noise on subsequent analyses, we apply Principal Component Analysis (PCA) to the data and keep enough dimensions to explain 80% of the total variance. Figure 3(a) illustrates OFC_*A*_(*t,v*) for all 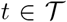 and 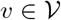 using t-SNE [van der Maaten and Hinton, 2008], a dimensionality reduction procedure that projects high-dimensional data onto a two-dimensional plane to facilitate visualization.

**Figure 3:**
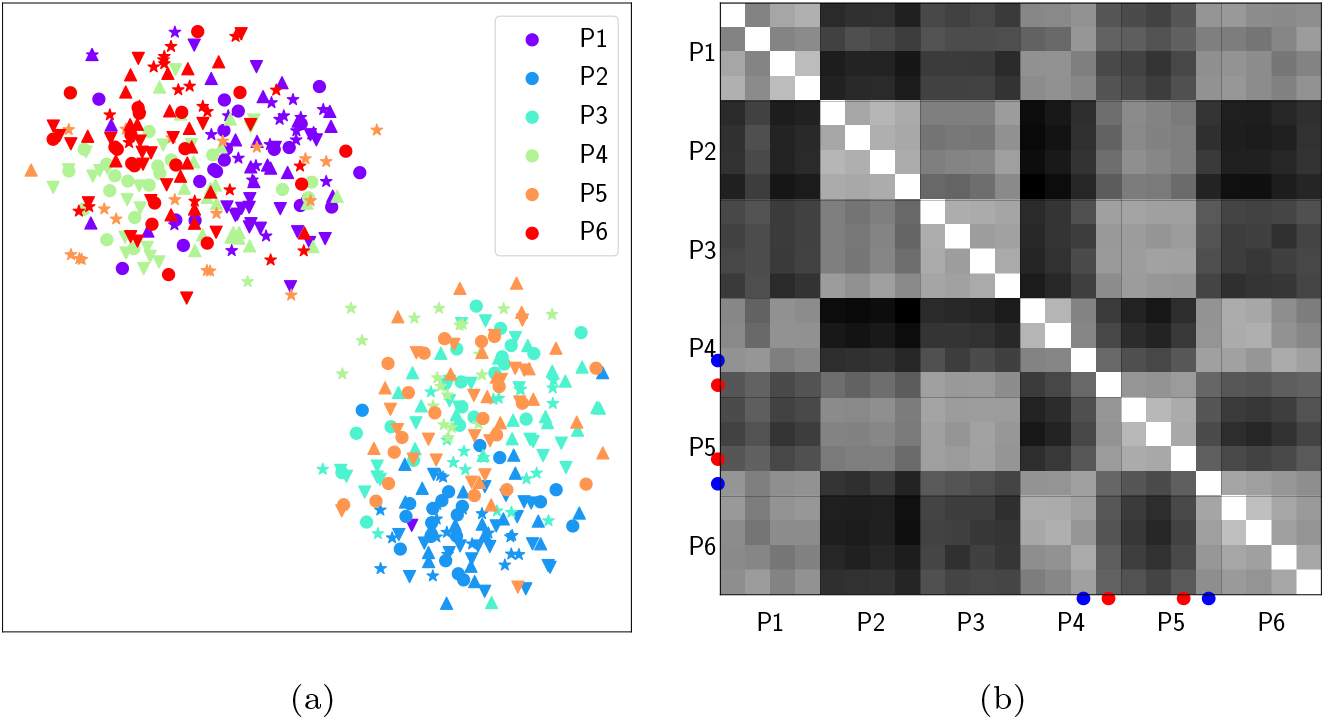
Visualization of sample representations and their representational dissimilarity matrix (RDM). Figure (a) plots the function OFCA(·,·) using t-SNE. There are a total of 480 points in this scatter plot, each of which is the OFC representation of a visit to a trial in problem *A.* Colors represent positions and the four shapes (circle, down triangle, up triangle, and star) correspond to the four sequences (S1a, S1b, S2a, S2b). We observe that, broadly, visits to rewarded trials appear closer to each other but farther from visits to non-rewarded trials. The heatmap in (b) is a rendering of the RDM computed from representations in (a). The rows and columns represent 24 unique trials, grouped by position and, within each group, sorted by sequence. Darker shades indicate a distance closer to 1. By construction, the RDM is symmetric with zeros on the main diagonal. We have marked positions P4 and P5 of sequences S2a and S2b with a blue (for rewarded) or red (for non-rewarded) filled disc to highlight the irregularity in S2b.

Given the setup of the odor sequence problem, the formulation above results in 480 distinct vectors representing as many *visits* that make up a single problem. We often find it useful to talk about the representation of a *trial* instead regardless of when that trial is visited. To facilitate such discussions and analyses, we define the representation of a trial *t* to be the mean of the 20 vectors that represent visits to *t*:

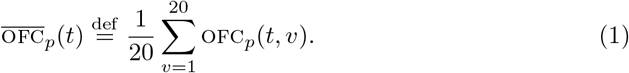

#### 3.1.2 Agent Neural Representations

Similar to OFC representations, we use 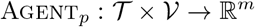 to express an agent’s mapping from visits of trials to *m*-dimensional representations. When the agent is A3cRnn, Agent_*p*_ is the output of the hidden layer of the RNN. For A3cFfn, it is instead the output of the last layer of its feed-forward network. Note that, unlike the biological subjects, agents’ output dimension, *m*, is constant across problems.

Mirroring once more how OFC representations are prepared, we train an agent 100 times from a random initial state and concatenate their individual representations to form Agent_*p*_(·,·):

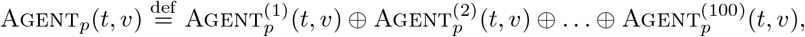

where 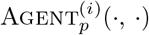 is the output of the *i*^th^ trained agent. We reduce the dimensionality of the raw representations using PCA to 32. Lastly, as with Equation (1), we define 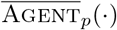 to produce the visit-independent representation of a trial.

#### 3.1.3 Geometry of Neural Representations via Representational Dissimilarity Matrices

As discussed in the preceding section, we view trials as vectors in a high-dimensional Euclidean space. But while that formalism is conceptually convenient, working with a potentially large set of vectors can quickly become cumbersome. We therefore seek to summarize an entire dataset of neural representations in a single object that captures some property of interest. One such property is the pairwise distance between vectors. That information begins to paint a picture of how vectors are arranged in a representational space, with larger distances taken to signal higher dissimilarity.

With that perspective, we define the distance between a pair of trials to be the angular distance between their respective representations. More precisely, if **u** and **v** are representations of two trials in ℝ^*n*^ (e.g., **u** and **v** may be the output of 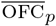 or 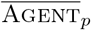), we record the following quantity as a measure of their distance:

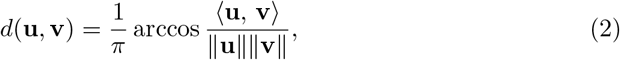

where 〈·, ·〉 denotes the inner product of two vectors, and ||·|| is the Euclidean norm. This distance is between 0 and 1, with a larger value indicating a higher degree of dissimilarity.

Using the pairwise distance between all trials in 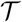, we construct a representational dissimilarity matrix (RDM) such as the one rendered in Figure 3(b). If *M* is an RDM, its (*i,j*)^th^ entry, *M_ij_*, records the distance as measured above between two trials, one corresponding to row *i* and another to column *j*. Naturally, *M* has zeros on its main diagonal as those entries reflect the distance between a trial and itself. Finally, by construction, an RDM is symmetric (i.e., *M*^T^ = *M* with *T* denoting the transpose operator).

RDMs appear in virtually every step of our study, illustrated and quantified. We plot RDMs to facilitate a visual inspection of patterns of dissimilarity among trials. In such illustrations we render the complete matrix as a heatmap. On the other hand, when quantifying RDMs it suffices to consider only the off-diagonal entries in the lower triangle (i.e., *M_ij_* for all *i* > *j* of an RDM *M*). That is because RDMs are symmetric by construction and, as a result, the lower half encapsulates all that is needed for analysis. We perform this masking transformation when answering quantitative questions.

### 3.2 Is Representational Geometry Independent of Problem Instance and Source?

We have just seen how an RDM summarizes a representational space. Specifically, an RDM captures the geometry of a dataset of neural representations by encapsulating pairwise distances between vectors. A natural question then is, given two datasets, possibly obtained from separate sources, can we compare the similarity of the structure of their respective spaces using RDMs? That constitutes our first question.

When investigating the inquiry above for the odor sequence problem, we use the following null hypothesis: Subjects and agents learn to represent all elements of the task structure but do not learn to delineate sequences. This entails that trial representations do not contain any information about sequences and, as such, are interchangeable within a single position. That means, for example, that swapping the representations of trials S1a3 and S2b3—both describing distinct trials in position P3—must, with great likelihood, lead to the same conclusion as the original sequence assignment.

In answering our first question under the null hypothesis above, we take the correlation between a reference RDM and a candidate RDM as the test statistic. More precisely, it suffices to compute Spearman’s rank correlation between the off-diagonal, lower-triangle entries of the two RDMs as a measure of the correlation between the two matrices, as done in previous studies [Popal et al., 2020]. Measuring rank correlation is justified because it is only of interest if the relative dissimilarity between trials is consistent across RDMs; crucially, we do not expect that trial dissimilarities correlate in absolute terms across problems.

Note that, when computing the correlation as explained above, it simply does not matter what data generated the RDMs; so long as datasets can be turned into RDMs we can easily compute the correlation between them. For example, in the context of the odor sequence problem, we can measure the correlation between an RDM generated by OFC recordings and another generated from an agent’s data.

Having computed our test statistic, we must now determine its significance. To that end, we use a permutation test to assess the likelihood that our test statistic is observed merely due to chance under the null regime. This test requires two steps. First, we transform the candidate RDM so as to eliminate any information about sequences. We do so by taking trial representations of each position and assigning them to a randomly selected sequence. In the resulting set, representations preserve all their original attributes, but are now presumed to have originated from a possibly different sequence—a valid configuration under the null regime. Figure 4 describes the procedure above pictorially. We then use this transformation to produce 100, 000 transformed RDMs and record their correlation with the reference RDM.

**Figure 4:**
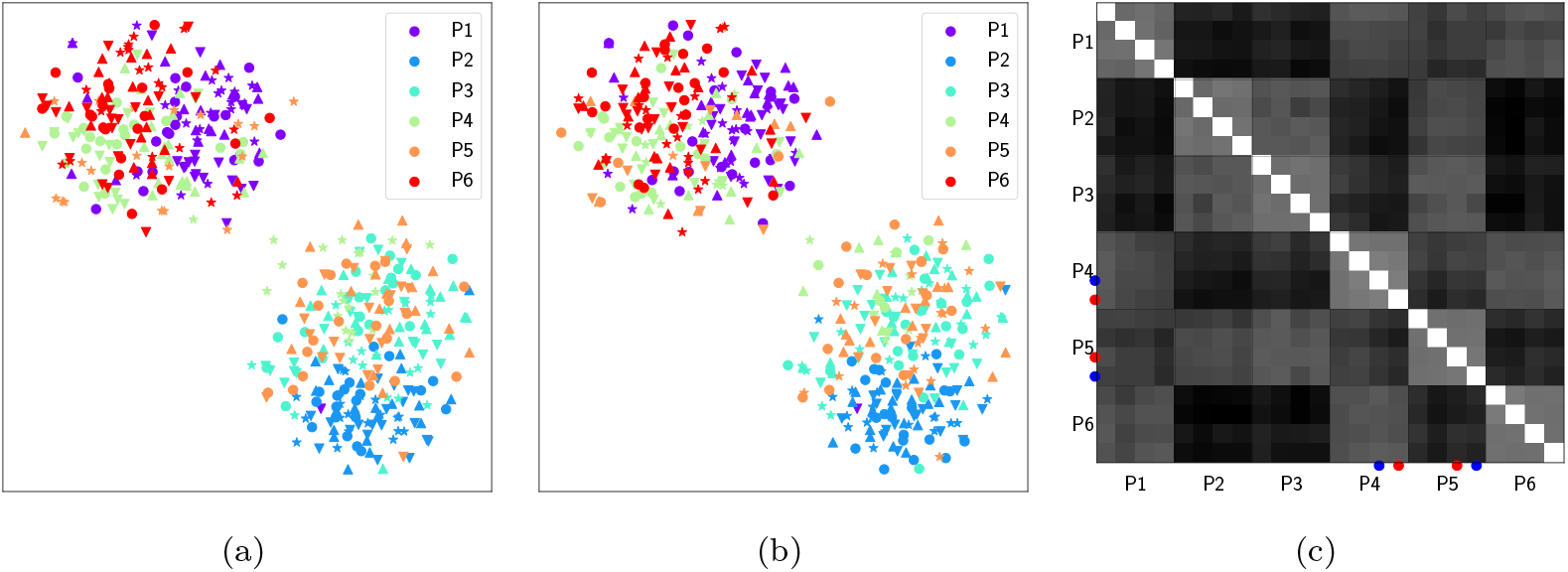
Visualization of a permutation under the null assumption and its resulting RDM. (a) is copied from Figure 3(a) depicts the original representations of 480 trial visits from problem A, where colors show positions and shapes sequences. (b) displays the same set of representations as in (a), but where, for every position, visits are assigned a new sequence that is chosen at random (i.e., colors stay the same but shapes change). (c) is the RDM resulting from permuted representations shown in (b).

The step above results in a distribution of correlations should the null hypothesis hold, referred to as the null distribution. In the second and final step, given the null distribution, we estimate the *p*-value of the test statistic as the number of permutations for which the test statistic exceeded the observed value plus 1, divided by the total number of permutations.

### 3.3 Are Geometrical Similarities Driven by Common Latent Factors?

The previous section described an approach to determine whether trials as represented in one space have a geometrical arrangement that is similar to those in another space. But simply because two structures look similar does not necessitate that the factors that give rise to the structure also correlate. For example, a set of vectors maintains the same structure when all vectors in the set are rotated by the same degree in the space, yielding perfect correlation between RDMs. In that sense, our method thus far is inconclusive, leading us to a second question: Is the similarity between two RDMs driven by correlated latent factors?

We answer the question above by investigating whether the structure of trials in a reference space can be accurately approximated from latent factors that are correlated between the reference and candidate spaces. For example, had we projected OFC representations of problem A (reference) onto the subspace that is correlated with problem *B* (candidate), would we observe similar pairwise distances between trials? In this section, we present a method to answer such questions.

The main difficulty in examining this new question lies in finding factors that are correlated between two spaces, as the reference and candidate spaces are assumed to be misaligned (e.g., have a different number of dimensions). In other words, there is no shared subspace that is “common” between the two spaces in the usual sense. To resolve that difficulty, we use Canonical Correlation Analysis (CCA). CCA is an optimization problem where, given two (misaligned) spaces *X* and *Y*, it finds subspaces *X*′ and *Y*′, where each latent feature in *X*′ (*Y*′) is a linear combination of features in *X*(*Y*), with the constraint that the *n*^th^ feature from *X*′ maximally correlate with the *n*^th^ feature from *Y*′. Throughout this work, with a slight abuse of terminology, whenever we speak of *the common space between two spaces*, we are referring to the correlated subspaces as determined by CCA. We use a regularized, generalized CCA [Bilenko and Gallant, 2016] in our work.

Taking this into account, our procedure consists of the following steps. First, we split the set of visits from the reference space into even and odd subsets, thereby forming disjoint training and test partitions. We then apply CCA to find a 10-dimensional common space between the space of the training set and a given candidate space. Using the resulting transformation, we project representations in the test set onto the latent space. Finally, we compute one RDM from the test set in the original reference space, and another from the projected test set in the latent space, and measure their correlation. We repeat this procedure by swapping the roles of even and odd visits. The mean of the two measurements becomes our test statistic. Figure 5 gives a schematic overview of this procedure.

**Figure 5:**
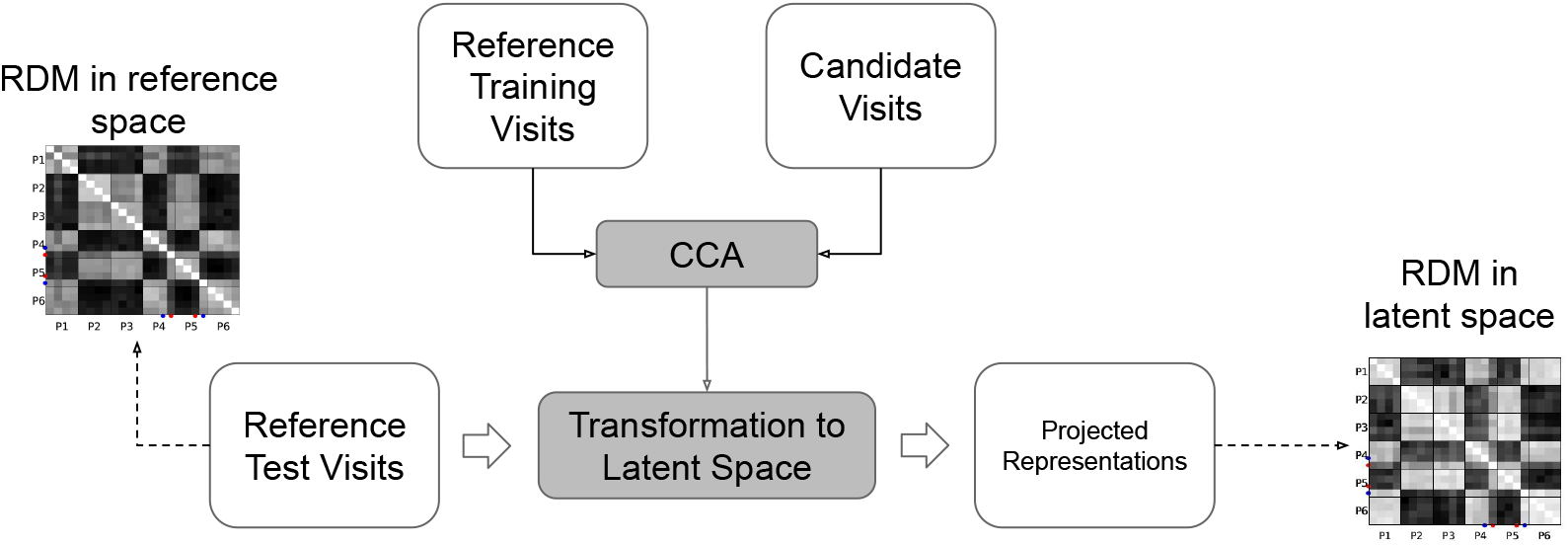
Diagram of a procedure to measure the contribution of common features to the geometric structure of trials. First, the reference representations are split into disjoint training and test visits. Given the training set and a candidate set of representations (e.g., from another problem or from an agent), Canonical Correlation Analysis (CCA) finds transformations into a lower-dimensional latent space. We then use the resulting transformation to project the test visits to the reference latent space. Finally, we measure the correlation between the RDM of the test set in the original space and the RDM of the test set in the latent space. We use the same procedure to obtain the null distribution but where the candidate representations are first permuted under the null assumption.

Having obtained a test statistic, we now turn to assessing its significance with respect to our null hypothesis using a permutation test. As required, we permute the candidate representations to obtain a new candidate space that is valid under the null assumption. With this new candidate, we follow the procedure above to calculate the correlation between RDMs of the original and projected test sets of the reference representations. By repeating this process 100, 000 times, we approximate the null distribution and estimate the *p*-value as described earlier.

### 3.4 Do Common Latent Factors Encode Similar Information?

The method in the preceding sections allow us to determine whether the geometry of trials is consistent between representational spaces and whether that consistency is due to common latent factors. But geometrical similarity and the existence of common latent factors do not imply that the two spaces encode similar information. It remains, then, to examine how informative the latent factors are in the following sense: Does having full knowledge of one representational space enable us to accurately deduce an optimal decision in another representational space? We develop a method to investigate that final question.

We formulate the question above as a decoding task in which, given the representation of a visit to a trial, the decoder predicts the optimal action. At a high level, we train a decoder on the representations from two randomly selected problems (training) and evaluate it on the remaining problems (test). While the test representations always come from the OFC, the training representations originate either from the OFC or agents: If from the OFC, we refer to the decoder as *d*_OFC_, otherwise we denote it by *d*_A3cRnn_ or *d*_A3cFfn_ depending on which agent provides the representations. We repeat the process above multiple times, each time choosing a different configuration of training and test problems, and report the mean and 95% confidence intervals.

One challenge in training a decoder that can work across spaces is the issue of misalignment, as touched on previously. We address that challenge as we did before using generalized CCA [Bilenko and Gallant, 2016]. We start with five sets, each consisting of 480 representations from a unique problem. Two of these sets are obtained from the OFC or the agent, depending on the type of decoder being trained, and the remaining three come from the OFC. From each of these sets, we leave out one visit to every trial. That leaves 456 representations in each set, with 24 held out for evaluation purposes. By applying CCA to these sets, we find 10-dimensional aligned latent spaces which we refer to as the common space.

Now that representations are in the common space, we take the two training problems to train a 5-Neighbors classifier. Given a test representation, such a classifier finds 5 representations closest to it—in the sense of Equation (2)—and returns the majority vote as the predicted action. Finally, we project the held-out representations from the test problems onto the common space, and evaluate the accuracy of the decoder on the projected set. This procedure is depicted in Figure 6.

**Figure 6:**
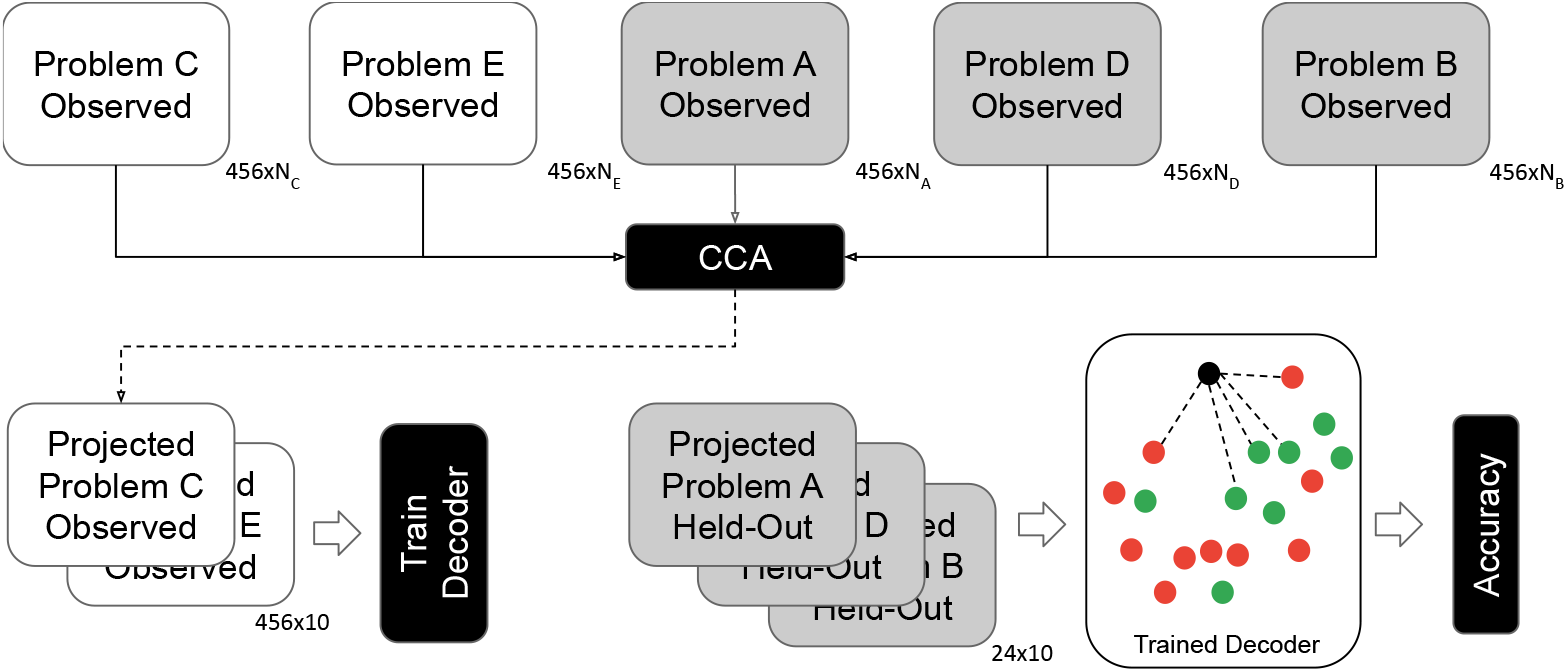
Diagram of the decoding method. **Top row**: We randomly divide the five problems into training and test classes—in this example, problems *C* and E are in the training class and the remaining three in test. In the case of *d*_OFC_, the representations for the training problems are obtained from the OFC, whereas in *d*_A3cRnn_ and *d*_A3cFnn_ they come from the respective agent. The representations for the test problems, on the other hand, always come from the OFC. From these sets, we leave out one visit per trial from every problem for evaluation purposes. Finally, we apply CCA to the (misaligned) spaces of the remaining representations to obtain 10-dimensional, correlated latent spaces. **Bottom row, left**: We project the representations from the training problems onto the latent spaces using the learnt CCA transformations, and train a decoder that predicts the optimal action given a representation. **Bottom row, right**: We project the held-out visits (24 in total) from the test problems onto the latent space found by CCA and measure the accuracy of the decoder on the resulting set. **Decoder** : We use a 5-Neighbors classifier. Given an unlabeled point (black), it finds 5 labeled points (green and red) from the training set that are closest to it (connected with dashed lines). The output of the decoder is the majority vote of these 5 points. In the illustrated example, the predicted label of the test point is green with 3 votes (60%).

## 4 Results

Having described our method, we now analyze the datasets of biological and artificial neural recordings from the odor sequence problem. We start by revisiting the key research question of Zhou et al. [2021]: Does OFC neuronal activity evolve to encode a representation that is common across different problem instances? We put this question in terms of the three inquiries of our methodology. We present the result of our analysis in the first subsection, confirming and extending the original findings.

We turn next to a direct comparison of the representations learnt by agents with neural recordings. As noted before, the behavior of the agents considered in this work is similar to the animals. Through our analysis, we wish to verify whether the neural codes generating those behaviors are also similar between the machine and the brain.

Our results focus primarily on showing that our methodology can be applied to neural recording data to achieve the goals we proposed, given a reasonable choice of agent. While we also show that our methodology can be used to compare models, we purposefully do not include any results on modelling behavior in isolation, as that question is already amply tackled in the literature (e.g., Song et al. [2017]).

### 4.1 Comparing OFC Activity between Problems

Our first results concern the neural activity in the subjects’ OFC only. As Zhou et al. [2021] found, OFC neuronal activity evolves to encode task features that are common to all problem instances. Here we re-examine their findings through the lens of the method developed in this work.

The first question asks whether the representational space learnt in one problem (say, *A*) is similar to those learnt in other problems, and if so, whether that similarity is significant. A negative result (i.e., insignificant correlation) would indicate that the data do not support the findings of Zhou et al. [2021] under the particular null hypothesis stated in Methodology, whereas a positive result clears the path for us to proceed to our second inquiry.

As discussed earlier, the OFC neural representations during Problem A can be summarized as an RDM, shown in Figure 7(a). The same can be done for every other problem, e.g. the RDM from Problem B is shown in 7(b). Finally, a sample RDM generated under the null hypothesis is shown in in 7(c). To answer our first question, we compute the correlation between a reference RDM (Problem *A*) and those of problems *B, C, D*, and *E*, shown as the blue bars in Figure 7(d). The figure also shows the mean correlation of each problem RDM with those of 100, 000 RDMs under the null distribution, and the resulting *p*-value. As is evident from this figure, the geometry of the representations learnt in the OFC appear to correlate significantly across problems.

**Figure 7:**
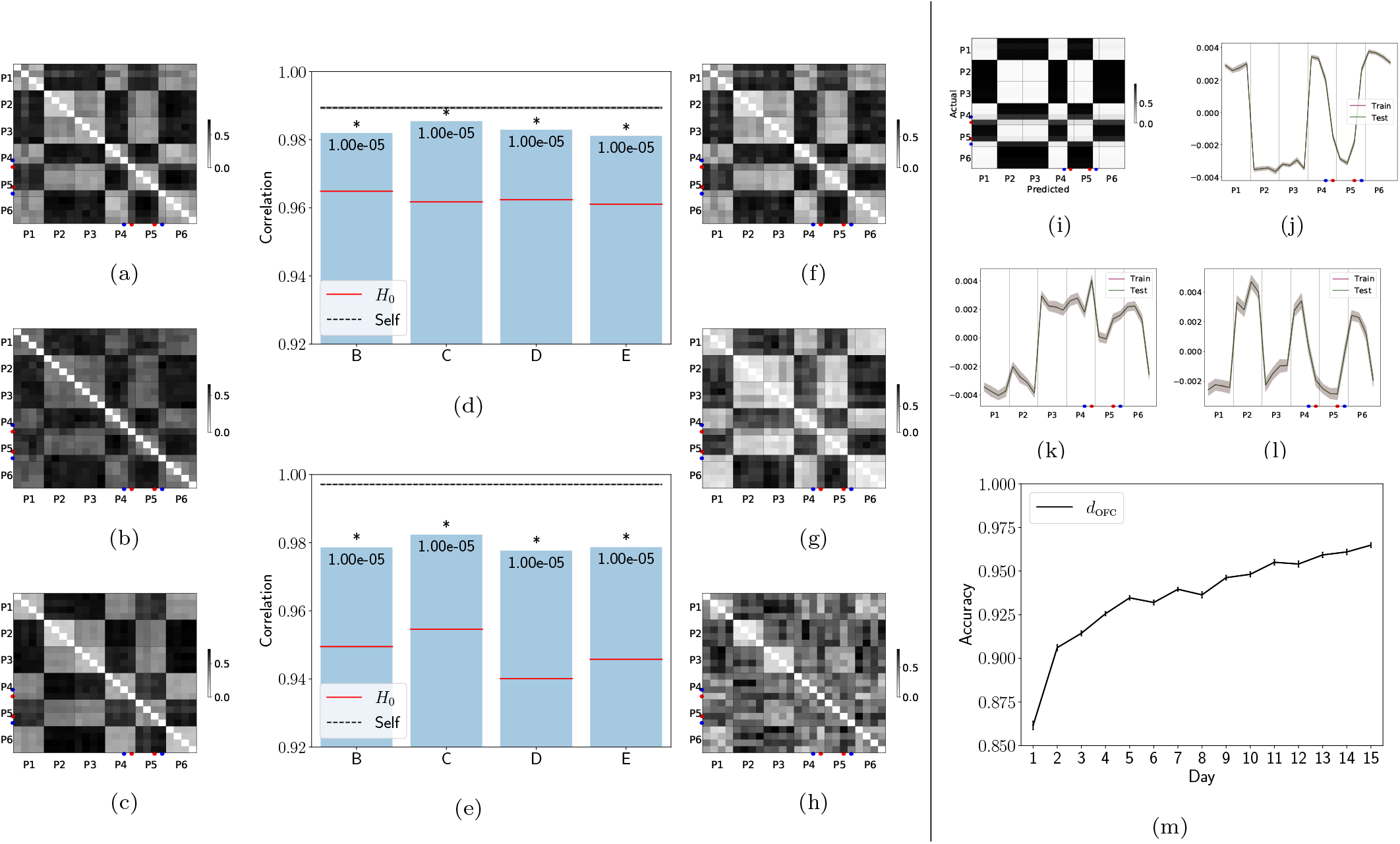
Examination of OFC representations from the final day of learning. The first column shows sample representational dissimilarity matrices (RDM) constructed from OFC representations for (a) Problem A, (b) Problem B, and (c) the null hypothesis, *H*_0_, that sequences in Problem B are indistinguishable. RDMs are partitioned by positions P1 through P6, with rows and columns within each partition corresponding to sequences S1a, S1b, S2a, and S2b. Darker shades indicate greater angular distance between representations of intersecting rows and columns. Figure (d) shows Spearman’s rank correlation between RDMs of Problem A with Problems B through E, along with the performance of Ho and noise ceiling (“Self”), the correlation of even visits of Problem A with odd visits. *p*-values noted in the figure are measured with a permutation test consisting of 100,000 permutations, with asterisk marking statistical significance (*p*-value < .05). Figure (e) similarly plots correlations, but where the reference is the test portion of Problem A and the object of comparison is a reconstruction of that same test set from the low-dimensional space that is common to Problem A and other problems, the null hypothesis, or itself. The third column visualizes sample RDMs generated from (f) the original representations for Problem A, and those reconstructed from the space common between Problem A and (g) Problem B and (h) the null hypothesis. The rightmost column presents the results of a decoding experiment where the objective is to determine the action (go/no-go) given the OFC representation of trials from one of three test problems, and where the decoder—denoted by *d*_OFC_—is trained using OFC representations of the remaining two problems. Figure (i) shows the optimal decision in each position within each of the four sequences. Figures (j), (k), and (l) are renderings of the first three latent factors from the train and test sets, which track task features (value, odor overlap, and positional alternation, respectively). Finally, (m) depicts the decoder’s accuracy over time, where data from each day of learning is used to train and evaluate a decoder for that day. Error bars show 95% confidence intervals.)

Now that we have established that representations exhibit similar geometrical properties across problems, we ask our second question: Are similarities driven by common latent factors? We apply the procedure illustrated in Figure 5 to this dataset and plot the results in Figure 7(e).

As shown in Figure 7(e), the RDM generated from representations in Problem A can be approximated, with a high degree of accuracy, by RDM reconstructions generated from latent factors that are in common with representations in other problems. We can see, for example, the similarity between an original RDM depicted in Figure 7(f) and its reconstruction from the common space in 7(g), which stands in contrast with 7(h), its reconstruction from the space that is in common with a null RDM.

We conclude, given the discussion above, that there exist common factors that drive the geometrical similarities observed earlier. But the existence of such factors alone does not reveal anything about the information they encode and, as such, is inconclusive. That brings us to our third and final inquiry: Does the information contained in the common space help us find the optimal policy in an unseen problem? We apply the decoding method to find out.

We train and evaluate the decoder *d*_OFC_, using the procedure explained in earlier sections, with the training and test set of representations both originating from OFC recordings. We repeat this exercise for all 15 days; that is, we train and evaluate *d*_OFC_ on data from day 1, day 2, through the final day, separately. Figure 7(m) plots the decoding accuracy over the course of learning, with the confusion matrix for the final day depicted in Figure 7(i). As a confirmation of the hypotheses of Zhou et al., additional days of learning brings about a higher cross-problem decoding accuracy. This indicates that, the common latent factors used by the decoder are not only similar geometrically, but that they also encode abstract information that enables generalization to other problems. Furthermore, continued learning leads to a refinement of such factors.

It is also instructive to take a look at the common latent factors between problems, which are used as input by the decoders. Figures 7(j), (k), and (l), respectively, display the first three factors shared between problems. Each of these orthogonal factors represents information for all six positions, across four sequences each. Interestingly, they correspond closely with the key aspects needed to perform the task: position value; uniqueness of odors to positions; and, positional alternation.

### 4.2 Comparing OFC Representations with A3cRnn

Whereas in the preceding section we studied generalization across problems in the OFC, in this section our main objective is to validate the A3cRnn model against the OFC recordings. To do that, we follow the same recipe as before and make similar inquiries into the space of representations.

While no modification to the method is necessary to facilitate this analysis, the datasets being compared now come from different sources. In the first inquiry, for example, we compare the geometrical similarity of the subjects’ space with the agent’s for each problem separately, as shown in Figure 8(d). For the second inquiry, as reflected in Figure 8(e), we measure how accurately a problem’s OFC RDM can be approximated from what it has in common with the agent’s representations. For each problem, we follow the procedure illustrated in Figure 5, using the OFC RDM as reference and the A3cRnn as the candidate. Finally, when training a decoder as part of the third inquiry, the training set is supplied by the agent while the test set is from the OFC—in other words, we are interested in the accuracy of a decoder trained on the agent’s representations when applied to the OFC representations. This follows the procedure illustrated in Figure 6, using agent’s representations of two problems as training data, and OFC representations of the remaining three problems as test data.

**Figure 8:**
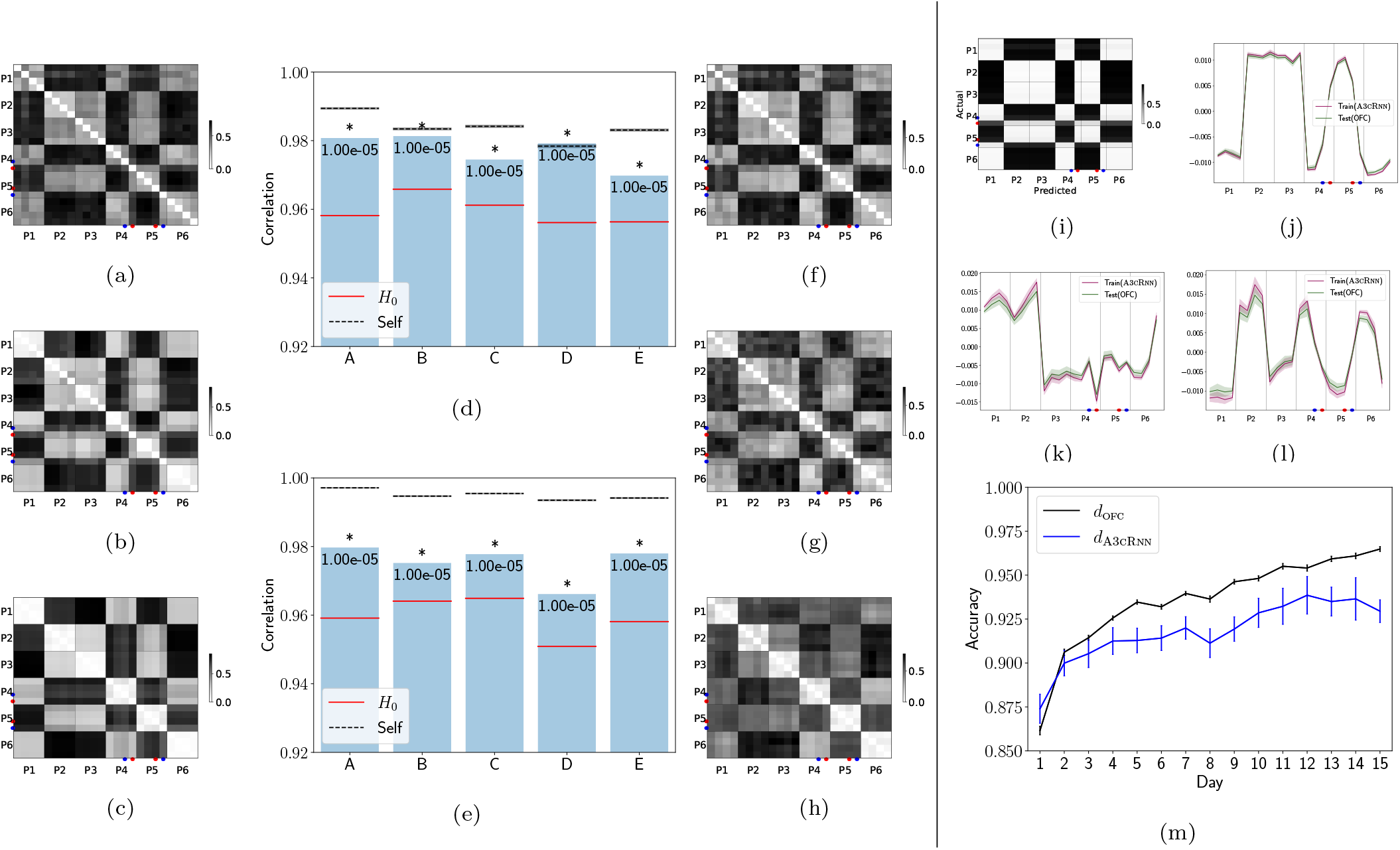
Examination the relationship between representations from OFC and A3cRnn on the final day or episode of learning. The first column shows sample RDMs constructed for Problem A from (a) OFC, (b) A3cRnn, and (c) the null hypothesis, Ho, that the agent does not differentiate between sequences. RDMs are partitioned by positions P1 through P6, with rows and columns within each partition corresponding to sequences S1a, S1b, S2a, and S2b. Darker shades indicate greater angular distance between representations of intersecting rows and columns. Figure (d) shows Spearman’s rank correlation between RDMs of OFC with A3cRnn, *H*_0_, and itself for Problems A through E. *p*-values noted in the figure are measured with a permutation test consisting of 100,000 permutations, with asterisk marking statistical significance (*p*-value < .05). Figure (e) similarly plots correlations, but where the reference is the test portion of the OFC dataset and the object of comparison is a reconstruction of that same test set from the low-dimensional space that is common to OFC and A3cRnn, the null hypothesis, or itself. The third column visualizes sample RDMs generated for Problem A from (f) the original OFC representations, and those reconstructed from the space common between OFC and (g) A3cRnn and (h) *H*_0_. The rightmost column presents the results of a decoding experiment where the objective is to determine the action (go/no-go) given the OFC representation of trials from one of three test problems, and where the decoder—denoted with *d*_A3cRnn_—is trained using A3cRnn representations of the remaining two problems. Figure (i) shows the optimal decision in each position within each of the four sequences, as determined by *d*_A3cRnn_. Figures (j), (k), and (l) are renderings of the first three latent factors from the train (agent) and test (OFC) sets, which track task features (value, odor overlap, and positional alternation, respectively). Finally, (m) depicts the accuracy of *d*_A3cRnn_ over time, where data from each day and its corresponding episode are used to train and evaluate a decoder. For reference, the figure also shows the performance of *d*_OFC_, a decoder that is trained and evaluated on OFC data only. Error bars show 95% confidence intervals.

Figure 8 shows the output of our analysis. These results are broadly consistent with what we observed in the preceding section. Remarkably, the A3cRnn agent has learnt representations that are significantly correlated with the neural recordings from the OFC, with common latent factors contributing significantly to structural similarities. Furthermore, decoding accuracy improves significantly as the subjects and agents go through their respective training. This suggests that, in addition to learning similar representations for the problem, experimental subjects and agents develop them in the same, gradual manner across the training regime. Finally, factors that maximally correlate between the OFC and agent spaces correspond with task features.

### 4.3 Comparing OFC Representations with A3cFfn

We have observed that A3cRnn learns representations that are similar to what is represented in the OFC in the context of the odor sequence problem. A3cRnn was designed to be a simple agent capable of learning the task and exhibiting a behavior across problems that is similar to the experimental subjects’ decisions. Our methodology does not require a minimal agent, and determining what a minimal agent might look like depends on the experimental task. However, we believe that one aspect of the model design—the presence of recurrent connections—is critical for performance, and our methodology provides an elegant way of demonstrating it. To analyze its contribution, we replace the recurrent neural network with a feed-forward one, resulting in the A3cFfn variant described in Artificial Agent.

Using the same procedure as before, we arrive at Figure 9. From the example RDMs rendered in this figure, we see that much of the geometrical structure is shared between the OFC and A3cFfn, and in fact, much of this structure can be accurately reconstructed from the common space. However, the representations for positions P4 and P5 in sequence S2 are misplaced in the representational space of A3cFfn. That seemingly minor mismatch leads to an effect that is detectable by our analysis: The correlation between RDMs, whether in the original form or after reconstruction, are not statistically significant for a majority of problems.

**Figure 9:**
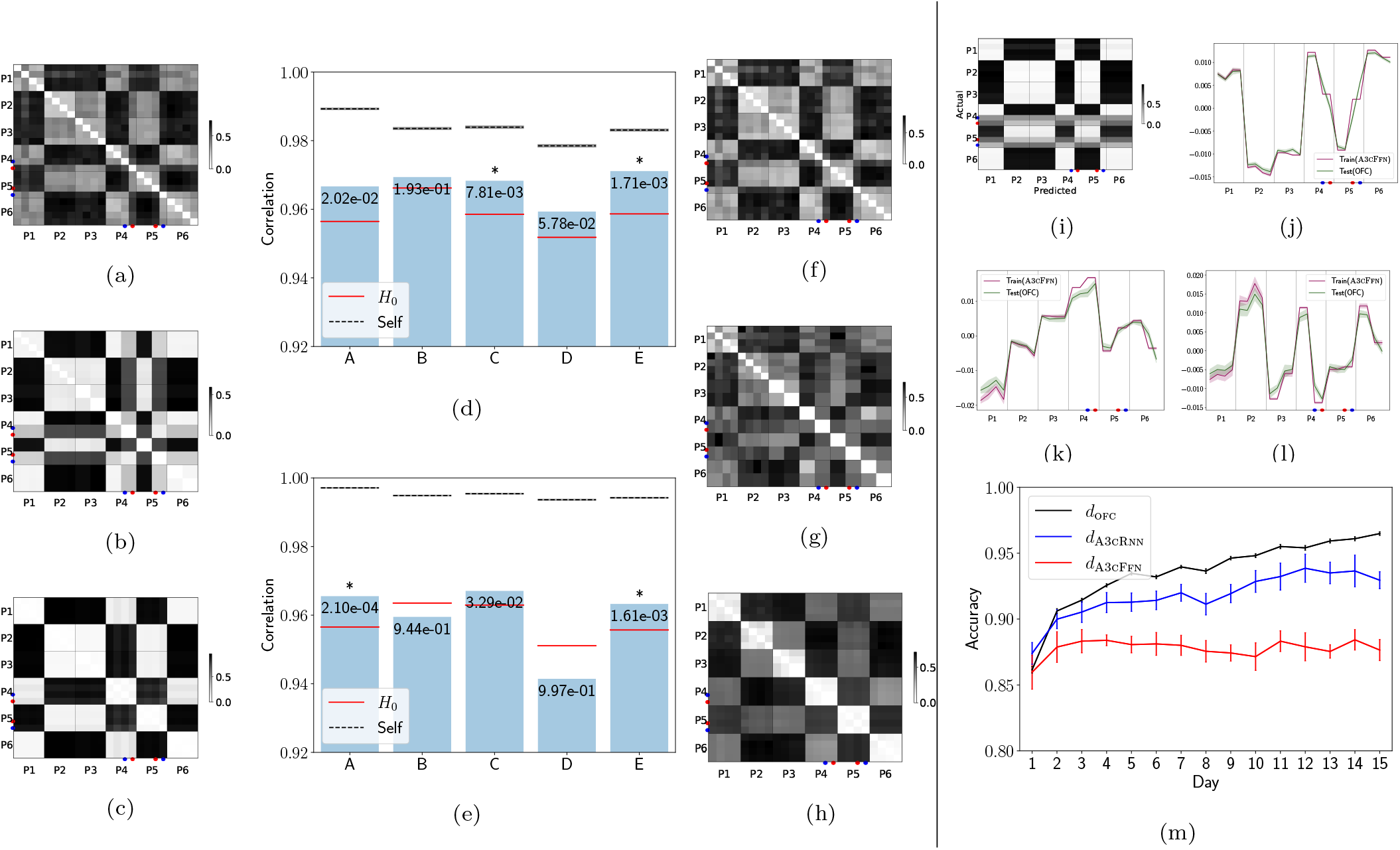
Examination the relationship between representations from OFC and A3cFfn on the final day or episode of learning. The first column shows sample RDMs constructed for Problem A from (a) OFC, (b) A3cFfn, and (c) the null hypothesis, *H*_0_, that the agent does not differentiate between sequences. A distant matrix is partitioned by positions P1 through P6, with rows and columns within each partition corresponding to sequences S1a, S1b, S2a, and S2b. Darker shades indicate greater angular distance between representations of intersecting rows and columns. Figure (d) shows Spearman’s rank correlation between RDMs of OFC with A3cFfn, *H*_0_, and itself for Problems A through E. *p*-values noted in the figure are measured with a permutation test consisting of 100,000 permutations, with asterisk marking statistical significance (*p*-value < .05). Figure (e) similarly plots correlations, but where the reference is the test portion of the OFC dataset and the object of comparison is a reconstruction of that same test set from the low-dimensional space that is common to OFC and A3cFfn, the null hypothesis, or itself. The third column visualizes sample RDMs generated for Problem A from (f) the original OFC representations, and those reconstructed from the space common between OFC and (g) A3cFfn and (h) *H*_0_. The rightmost column presents the results of a decoding experiment where the objective is to determine the action (go/no-go) given the OFC representation of trials from one of three test problems, and where the decoder—denoted with *d*_A3cFfn_—is trained using A3cFfn representations of the remaining two problems. Figure (i) shows the optimal decision in each position within each of the four sequences, as determined by *d*_A3cFfn_. Figures (j), (k), and (l) are renderings of the first three latent factors from the train (agent) and test (OFC) sets. Finally, (m) depicts the accuracy of *d*_A3cFfn_ over time, where data from each day and its corresponding episode are used to train and evaluate a decoder. For reference, the figure also shows the performance of *d*_OFC_, a decoder that is trained and evaluated on OFC data only, and *d*_A3cFfn_, a decoder that is trained on data from A3cRnn. Error bars show 95% confidence intervals.

Perhaps more interestingly, a decoder trained on representations from A3cFfn and evaluated on the OFC does significantly poorly compared to *d*_OFC_ and *d*_A3cFfn_. Moreover, its accuracy plateaus immediately after the first episode of training, suggesting that the agent quickly learns the representation of a default sequence, but appears unable to learn an optimal policy for the reversals.

A look at the factors shared between the two spaces reveals no obvious correspondence with task features, and in some cases, highlights what the agent learns incorrectly. For example, Figure 9(j) captures the value of each trial correctly with the notable exception of the reversals (i.e., positions P4 and P5 in S2). Figure 9(k) does not represent any of the task features, and, while Figure 9(l) appears to encode positional alternation, it again fails to correctly capture the reversals.

We compare A3cRnn and A3cFfn directly in terms of their performance on the second inquiry: The accuracy of the reconstruction of representational space of the OFC from what it has in common with one agent versus the other. This is plotted in Figure 10. It is clear that A3cRnn performs significantly better than A3cFfn on all problems.

**Figure 10:**
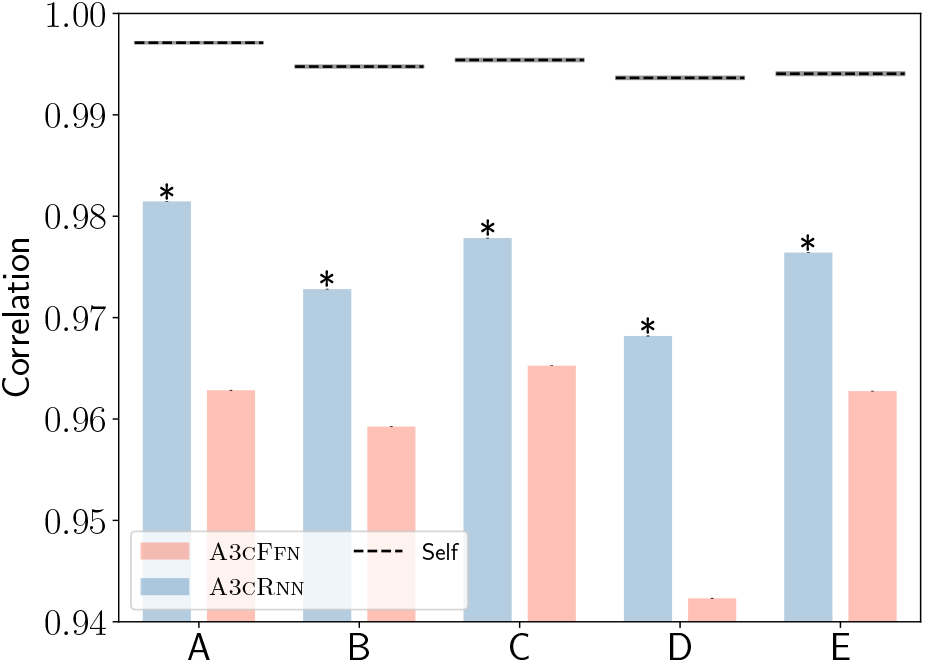
A comparison of A3cRnn and A3cFfn. For each problem, the OFC representations are split into train and test subsets. The train portion is aligned with a target representational space (either self, A3cRnn, and A3cFfn), and the resulting transformation is then applied to the test portion. The correlation between the RDM of the original and transformed test set are reported in the figure, with asterisks marking statistical significance (*p*-value < .05).

## 5 Discussion

We began this work with a simple observation: The computational neuroscience literature increasingly utilizes Deep RL to explain complex behavioral data. Seeing as the strength of Deep Neural Networks rests in their ability to learn rich latent representations, we argued that validating a Deep RL model must include an examination of the representational space, not just behavior. This is especially important in neuroscientific studies, where the neural mechanisms that cause a particular behavior have a higher scientific value. Crucially then, just as we validate the behavior of an artificial agent against real behavioral observations, we must compare neural codes learnt by a Deep RL agent with neural recordings obtained from a biological subject as the two navigate and learn the same task.

On the basis of the arguments above, we proposed a methodology to test hypotheses about representational spaces learnt by artificial agents as a combination of three key questions. The first question tests whether the representational similarity between two spaces is higher than what might be expected by chance, measured against a stringent null regime. The basic idea of comparing the geometry of the spaces of neural representation against those of a reference artificial agent or human subject is not novel and is known as Representational Similarity Analysis [Kriegeskorte et al., 2008] or RSA. However, if the similarity is significant, it allows the posing of a second question: Is this similarity driven by factors that are shared between the two spaces, or by factors that are entirely different—a question that RSA does not answer. In extending RSA to the second question, we use a cross-validated, non-parametric statistical procedure that is based on Canonical Correlation Analysis of neural representations, as illustrated in Figure 5.

Finally, the third question asks whether the common latent factors encode similar information. To answer, we test whether a decoder trained to predict the optimal action from latent factors in one domain (e.g. artificial agent) will do so accurately when tested on latent factors in another domain (e.g. experimental subject). We introduce a non-parametric statistical procedure combining Canonical Correlation Analysis and a classifier to carry this test, illustrated in Figure 6.

The questions and techniques we introduce are applicable to any reinforcement learning task that can be characterized as a Markov Decision Process (MDP), thereby covering a broad range of neuroscientific experiments, in particular, those described in Song et al. [2017]. This methodology is thus a stepping stone towards validating Deep RL models of behavior using neural data.

We demonstrated the utility of our method by re-analyzing a recently published study of transfer learning between several odor sequence tasks. We trained Deep RL agents to solve these tasks, which had hitherto not been accomplished, and used our methodology to show that they develop a representational space that contains similar information as the animal subjects. This study of representations can be pursued at any level of granularity (e.g., the individual events making up a trial).

Perhaps more interestingly, our method can help determine the necessity or role of individual components in the architecture of a Deep RL agent, and the appropriateness of their hyperparameters, even if a modification of that nature leads to a small effect. For example, after removing a recurrent mechanism from our default artificial agent, it became apparent that the resulting representational space deviated significantly from the biological space. Even though the behavioral effect was limited to two trials (out of 24), the deformation of the geometrical space was enough to quickly establish the importance of temporal signals in solving the task.

We believe that, with the rise in interest in Deep RL agents and their integration into computational models of natural phenomena in neuroscience [Botvinick et al., 2020], our proposal for a methodology to validate agents with neural data is warranted and, in fact, necessary. The method presented here is applicable to any Deep RL agent that learns an MDP; so long as there are neural representations to be examined, our method can be readily used. This generality enables researchers to assess the suitability of more advanced Deep Learning modules or map hierarchies of representations to different parts of a circuit in the brain, and help shed light on their function.

## 6 Acknowledgements

This work was supported by the National Institute of Mental Health Intramural Research Program (ZIC-MH002968, S.B., P.M., and F.P.) and the National Institute on Drug Abuse Intramural Research Program (ZIA-DA000587, G.S. and K99DA049888, J.Z.).

